# ROS accumulation and associated cell death mediates susceptibility to *Alternaria brassicae* in *Arabidopsis* accessions

**DOI:** 10.1101/581025

**Authors:** Sayanti Mandal, Sivasubramanian Rajarammohan, Jagreet Kaur

**Affiliations:** Department of Genetics, University of Delhi, South Campus, Benito Juarez Road New Delhi -110021, India; Current Address - National Agri-Food Biotechnology Institute, Sector-81, Mohali, Punjab - 140306, India

**Keywords:** ROS, cell death, *Alternaria brassicae*, RBOH, Jasmonic acid

## Abstract

*Alternaria brassicae* is a necrotrophic fungal pathogen capable of infecting most of the agriculturally important *Brassica* species. The mechanisms underlying invasion of *A. brassicae* and host responses are unknown. In the present study, we exploited the natural variation in *Arabidopsis* to understand the molecular and cellular mechanisms underlying resistance to *A. brassicae*. Using a subset of resistant (Ei-2, Ull2-3, Lz-0, and Cvi-0) and susceptible (Gre-0, Est-1, and Zdr1) accessions, we show that the susceptibility to *A. brassicae* is associated with higher ROS accumulation and cell death. Susceptibility to *A. brassicae* was reduced in the *rboh* (*D, E* and *F*) mutants that are incapable of producing ROS, suggesting that *RBOH D, E and F* may act as negative regulators of defence against this pathogen. Additionally, our data also supports the hypothesis that the Jasmonic acid (JA), Ethylene (ET) and Abscisic acid (ABA) signalling pathways positively contribute to resistance against necrotrophic pathogens. In summary, these results reveal the central role of ROS and cell death in the pathogenesis of *A. brassicae* and expand our understanding of plant-necrotroph interactions.

## Introduction

Necrotrophic pathogens actively kill host tissues as they invade the host and obtain their nutrients from the dead tissues/cells. Pathogenesis of necrotrophs usually involves extensive necrosis and tissue maceration. This is in stark contrast to the biotrophic pathogens, which derive their nutrients from living host tissues. The infection processes, nature of secretory proteins and associated host defence responses vary significantly between biotrophs and necrotrophs. One of the key differences is how cell death in the host affects the pathogenesis of biotrophs and necrotrophs.

Cell death or Hypersensitive Reaction (HR)-induced cell death in the host effectively stops biotrophic infection and is considered a typical resistance response of the host plant species. Cell death confines biotrophs by limiting or cutting off the nutrient supply and restricting the pathogen growth. However, cell death can be successfully used by necrotrophs to proliferate within the host. Activation of cell death pathways has been shown to promote susceptibility to broad range necrotrophs such as *Botrytis cinerea* and *Sclerotinia sclerotiorum* (Govrin & Levine 2002; Ranjan *et al.*, 2017; van Baarlen *et al.*, 2007).

A typical hypersensitive reaction is initiated by the generation of Reactive Oxygen Species (ROS) followed by localised cell death. Besides defense, ROS is produced by plants as by-products of many key processes such as respiration, primary metabolism, photosynthesis, and responses to abiotic stresses. Rapid production of ROS or oxidative burst is one of the earliest responses of plants to pathogen attacks. Various studies have shown the involvement of ROS in cell death (Van Breusegem & Dat 2006). H_2_O_2_, a versatile reactive oxygen species, has multiple functions in plant-pathogen interactions. In addition to a direct antimicrobial effect, it also triggers cell wall cross-linking, induces resistance gene expression and hypersensitive response (Shetty *et al.*, 2008). The rapid accumulation of ROS, cell death and callose deposition correlate with disease resistance in many biotrophic and hemibiotrophic pathosystems (Marino *et al.*, 2012; Rate & Greenberg 2001). The necrotrophic pathogens, in contrast, thrive on the cell death caused by excessive ROS production during the recognition phase. Earlier studies have shown the importance of pathogen-responsive host H_2_O_2_ in promoting cell death in the host thereby causing expansion of disease lesions to facilitate necrotrophic fungal infection (Govrin & Levine 2000). Williams *et al.*, (2011) showed that oxalic acid produced by *S. sclerotiorum* induces host ROS in compatible interactions. The role of ROS in facilitating infection is therefore seemingly contradictory and depends on the pathogen’s lifestyle. However, the mechanisms involving the spatiotemporal control of ROS metabolism during plant-pathogen interactions are largely unknown.

Resistance to necrotrophic pathogens is based on defence responses regulated by jasmonic acid (JA) and ethylene (ET) signalling pathways (Glazebrook 2005), production of antimicrobial metabolites (Ferrari *et al.*, 2003; Glawischnig *et al.*, 2004), scavenging ROS (Wolpert *et al.*, 2002), and control of cell death (Govrin & Levine 2002). However, the current paradigms of the role of ROS-induced cell death in plant-necrotroph interactions are largely derived from studies on archetypical broad host range necrotrophic pathogens such as *B. cinerea* and *S. sclerotiorum*. Whether these paradigms extend to all necrotrophic pathogens is still unknown.

*Alternaria brassicae* is a necrotrophic pathogen with an intermediate host range, infecting almost all species within the *Brassicaceae* family and causes Black leaf spot, a major fungal disease in oilseed mustard. In the absence of any strong genetic resistance identified so far in the Brassicas, insight into the plant-pathogen interactions and host resistance mechanism are very limited. Recently, we established the *Arabidopsis-A. brassicae* pathosystem in our lab. The interaction and disease responses in the model pathosystem appear to be closely resembling the natural host *Brassica juncea*-*A. brassicae* interaction (Mandal *et al.*, 2018). Variation in the response of a set of natural *Arabidopsis* accessions to *A. brassicae* was observed. Furthermore, conventional and association mapping approaches revealed that the variation in resistance to *A. brassicae* in *Arabidopsis* accessions was quantitative and possibly orchestrated by multiple genes (Rajarammohan *et al.*, 2017; Rajarammohan *et al.*, 2018). The molecular mechanism by which the accessions resist infection by *A. brassicae* is still elusive. Therefore, the main goal of this work was to study the molecular and cellular mechanisms underpinning *Arabidopsis*-*A. brassicae* interaction by exploiting the subset of resistant and susceptible accessions available to us. We show that natural variation in resistance to *A. brassicae*, a necrotrophic pathogen, correlates to differential ROS accumulation and associated cell death. Additionally, the phytohormones viz. JA, ET and ABA (abscisic acid) were found to positively contribute to resistance.

## Methods

### 1. Plant materials and growth conditions

Seeds of *Arabidopsis thaliana* accessions were obtained from the Nottingham *Arabidopsis* Stock Centre (NASC): Cvi-0, Ei-2, Est-1, Gre-0, Lz-0, Ull2-3, and Zdr-1. The mutants *rbohD* (CS9555), *rbohE* (SALK_146126C), *rbohF* (CS9557) and Col-0 (Wild type) were obtained from Arabidopsis Biological Resource Centre (ABRC). The mutants were confirmed homozygous lines, however we independently confirmed the homozygosity of the mutants (Supplementary Fig. 4). Seeds were surface sterilised, stratified at 4 °C in the dark for two days, and plated on Murashige Skoog (Titan Media) for ten days under short day condition (10 h light/14 h dark). Ten-day old seedlings were thereafter transferred to 3-inch pots containing autoclaved soilrite mix (1 soilrite: 1 vermiculite: 1 perlite) and grown under short-day conditions at 22 °C and 60-80% relative humidity. 5-6 weeks old plants having approximately 8-10 well expanded true leaves were used for infection.

### 2. *Alternaria brassicae* growth condition and infection assay

The *A. brassicae* J_3_ isolate used was originally isolated from the infected *B. juncea* grown in Delhi, India, purified by single spore isolation, and maintained on Radish Root Sucrose Agar (RRSA, pH 7.0) medium at 22 °C under a 12 h light/dark cycle. For plant infections, conidia were harvested from two-week-old *A. brassicae* cultures. Spore concentration was adjusted to 10^3^-10^4^ conidia ml^−1^. The plants were infected using drop inoculation method by placing four drops of 5 µl spore suspension on each leaf. Six to eight leaves were infected per plant. Inoculated plants were covered with perspex domes and placed in infection chambers to ensure 90% humidity under short day condition. Infection phenotypes were assessed, based on number of necrotic lesions formed, using a previously described Disease Index (DI) score (Rajarammohan *et al.*, 2017). The samples for microscopic analysis were collected at 24, 48, 72 and 96 hours post infection (hpi).

### 3. Hydrogen peroxide (H_2_O_2_) detection by 3-3’ Diaminobenzidine (DAB) staining

Detection of H_2_O_2_ was performed using a protocol modified from Thordal-Christensen *et al.*, (1997). Briefly, infected whole leaf samples were collected at different time points post inoculation and placed in a 24-well microtiter plate filled with 400-600 μl of 3-3’Diaminobenzidine (DAB) solution (1 mg/ml in water). The volume of the DAB solution used was adjusted to ensure that leaves were completely immersed. A gentle vacuum was applied to ensure that DAB infiltrated into the leaves. The plates were covered with aluminium foil and placed in an incubator at 22 °C for three hours. Following the incubation period, the leaves were transferred into 50 ml centrifuge tube and de-stained in ethanol: acetic acid solution [3:1 (v/v)] overnight.

To calculate the area of H_2_O_2_ spread, 96 hpi stained leaf samples were observed under Olympus SZX10, and the area measured using the inbuilt program ProgRes® capture Pro2.7 JENOTPIK Laser. Collected image data sets of *rboh*D, E and F mutants and Col-0 (WT) were subsequently analysed with the digital image analysis programs ImageJ for DAB staining (H_2_O_2_ accumulation).

### 4. Trypan blue staining to determine fungal growth and detection of cell death

To evaluate the extent of fungal spread, penetration structures and plant cell death, pathogen-infected leaves were stained with lactophenol-trypan blue. Infected leaf samples were collected in 50 ml tubes, at different times post inoculation and an adequate volume of lactophenol-trypan blue solution (equal volume of lactic acid: glycerol: phenol: water) was added. The tubes were placed in a boiling water bath for one minute, and then the staining solution was replaced with chloral hydrate (1 g/ml) for clearing the samples (Bartsch *et al.*, 2006). Chloral hydrate solution was changed twice every 24 hours.

The area of cell death was analysed using the same methodology as carried out above for calculating area of H_2_O_2_ spread.

### 5. RNA isolation and transcript analysis

Total RNA was extracted from 6-8 leaves collected from six plants of each genotype in each experiment using the RNeasy plant mini kit according to the manufacturer’s recommendation (Qiagen, Gaithersburg, MD, U.S.A.). First-strand cDNA was synthesised from 1 µg of total RNA using MMLV Reverse Transcriptase 1st-Strand cDNA Synthesis Kit (Epicentre Biotechnologies, Madison, USA) as per manufacturer’s protocol. PCRs were carried out using the standard setting in a QuantStudio 6 Flex Real-Time PCR System (ThermoFisher Scientific, Waltham, MA, U.S.A.). The mean cycle threshold (Ct) values of the genes were calculated from replicate wells. The mean Ct values of the target genes were normalised to the Ct values of the endogenous controls (TIP41-like gene-At4g34270 and UBC8-At5g41700) for each sample. Further, to calculate the relative expression levels, the dCt values of the infected samples were normalised to the mock (distilled water) inoculated samples of each time point. These values from the three individual biological experiments were subjected to data standardisation and compared using a described method (Willems *et al.*, 2008). Sequences of primers used in qPCR have been listed in Supplementary Table S5.

## Results

### 1. Differential response of *Arabidopsis* accessions to *Alternaria brassicae* infection

Rajarammohan *et al*., 2017 reported a huge variation in the response of different *Arabidopsis* accessions to *Alternaria brassicae* infection, varying from highly susceptible to highly resistant. To understand the cellular and molecular basis of the observed differential responses, a subset of resistant (Ei-2, Ull2-3, Lz-0 and Cvi-0) and susceptible (Zdr-1, Gre-0 and Est-1) accessions were selected for detailed analysis. Although the Disease Index (DI) for each of the susceptible accessions, calculated based on the number of necrotic lesions developed per leaf, was similar, the lesion growth rate in each accession showed considerable differences. In Zdr-1, the highly susceptible accession, the lesions spread to an average diameter of 0.41 cm and were associated with chlorotic halos around them. By 7 dpi, the lesions expanded and coalesced to cover the entire leaf surface resulting in a total collapse of the infected leaves. In other susceptible accessions, Est-1 and Gre-0, necrotic lesions were associated with chlorosis but did not expand enough to coalesce at the end of 7 days. As shown in Fig. 1, the susceptible accession develops spreading lesions with a size ranging from 0.33-0.41cm. In contrast, the resistant accessions Ei-2, Ull2-3, Lz-0 and Cv-8 had a lower DI, with only 1-2 infection sites developing into pinpoint (limited) lesions by 7dpi (Table 1 & Fig. 1). Although the disease was scored at 7 dpi, water-soaked lesions start appearing in the accessions as early as 3 dpi. Therefore, the initial time points of 24 hpi, 48 hpi, 72 hpi, and 96 hpi were used for the microscopic analysis of host responses.

**Table 1.** Disease phenotypes of different Arabidopsis accessions after infection with *Alternaria brassicae* and scored at 7 days post infection (dpi)

| S.No. | Accessions | Interaction phenotype | Disease Index <sup>α</sup> | Lesion size <sup>β</sup> (cm) |
| --- | --- | --- | --- | --- |
| 1. | Zdr-1 | Susceptible | 89.96 ± 6.13 | 0.41 ± 0.07 |
| 2. | Est-1 | Susceptible | 89.52 ± 4.50 | 0.36 ± 0.06 |
| 3. | Gre-0 | Susceptible | 79.42 ± 11.72 | 0.33 ± 0.04 |
| 4. | Ull2-3 | Resistant | 19.17 ± 4.79 | 0.03 ± 0.01 |
| 5. | Lz-0 | Resistant | 14.51 ± 5.43 | 0.03 ± 0.01 |
| 6. | Cv8 | Resistant | 10.99 ± 7.63 | 0.02 ± 0.01 |
| 7. | Ei-2 | Resistant | 4.45 ± 4.80 | 0.03 ± 0.01 |

<sup>α</sup> - Data are mean ± S.D., calculated from three independent experiments.
<sup>β</sup> - Data are mean ± S.D., calculated from three independent experiments.

**Fig. 1.**
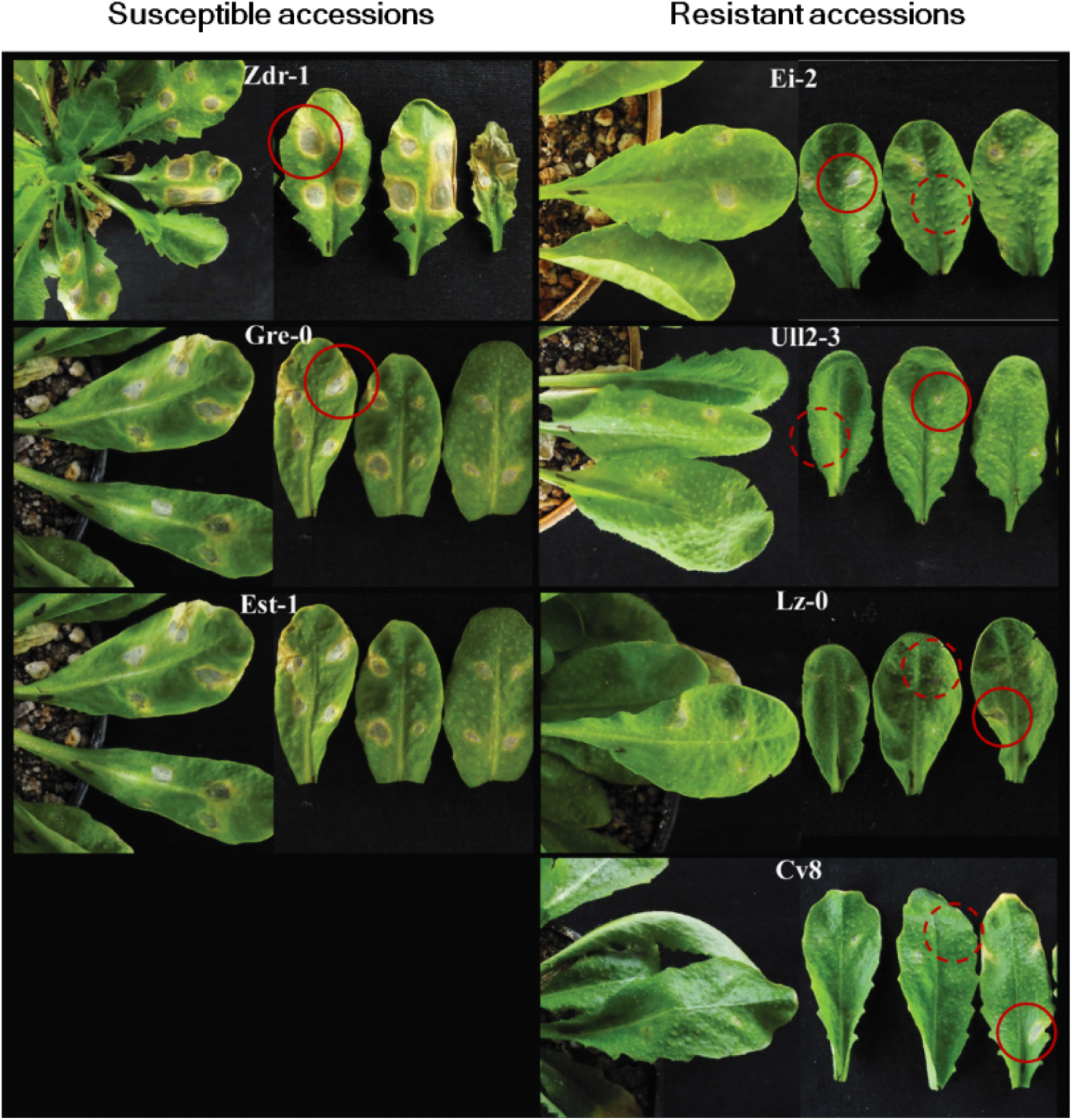
*Alternaria brassicae* infection phenotype on selected *Arabidopsis* accessions (7dpi). Left panel: Susceptible accessions Zdr-1, Gre-0, and Est-1 showing enlarged lesions at the site of inoculation and Right panel: Resistant accessions Ei-2, Lz-0, Ull2-3, and Cv8 with restricted lesions. Circles show the necrotic lesion, dotted circles on resistant accessions are regions of inoculation where microscopic cell death has been elicited

### 2. Comparable spore germination and penetration on resistant and susceptible accessions

Chemical and physical barriers at the host surface form the first layer of defence and could either inhibit or delay spore germination or fungal penetration. To determine the role (if any) of the first layer of defence in disease resistance against *A. brassicae* we compared spore germination efficiency and frequency of penetration by *A. brassicae* on both resistant and susceptible accessions. No significant difference was observed in the spore germination or fungal penetration efficiency in the two contrasting subsets. Spore germination initiated on both the susceptible and resistant accessions within 6 hpi (Fig. 2a & a’; Fig. 2b & b’) and by 24 hpi more than 90% of the spores had germinated. All the cells in the conidiophore were equally competent to germinate and, in several instances, multiple (3-4) hyphae emerged from the same conidia (Fig. 2a). At 24 hpi multiple, thick-branched hyphae showing dichotomous branching pattern were observed over the host surface (Fig. 2c & c’). Over multiple experiments, the spore germination efficiency was >99% on both resistant and susceptible accessions (Supplementary Table S1).

**Fig. 2.**
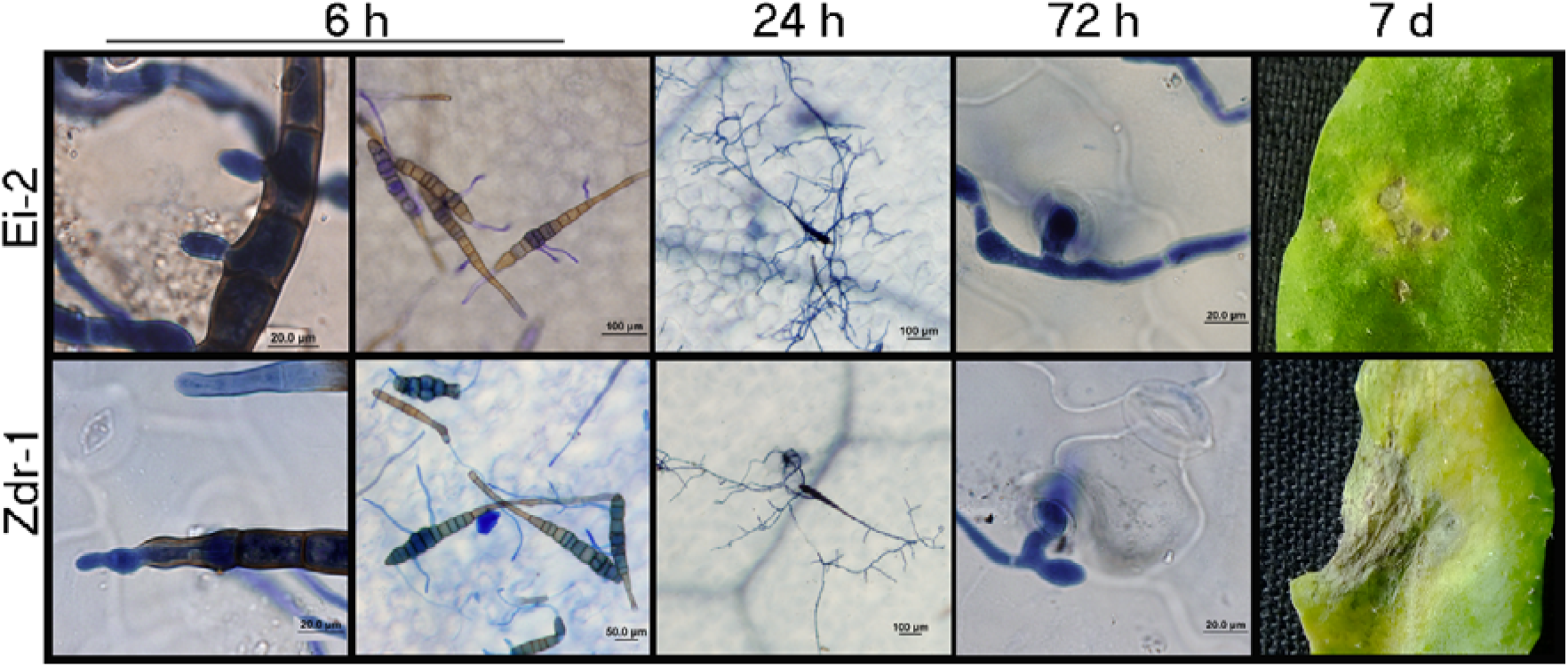
Comparative spore germination, hyphal growth, penetration and macroscopic infection phenotype on resistant (Ei-2) and susceptible (Zdr-1) *Arabidopsis* accessions upon *Alternaria brassicae* infection. (a and a’) Formation of germ-tubeat 6 hpi in Ei-2 and Zdr-1 (b and b’) Germinating *A. brassicae* spores at 6 hpi. (c and c’) Conidia producing 3-4 branched hyphae spreading over the surface of Ei-2 and Zdr-1 at 24 hpi. (d and d’) Appressoria like structure penetration through stomata on the surface of Ei-2 and Zdr-1 at 72 hpi. (d and d’) Phenotypic differences in the development of necrosis in resistant Ei-2 and susceptible Zdr-1

Following germination, *A. brassicae* invades the host tissue by developing **<u>A</u>**ppressoria **<u>L</u>**ike swollen, globular **<u>S</u>**tructures (ALS) at the hyphal apices. Similar Appressoria like structures were also observed when the fungus invaded its natural host *B. juncea* (Mandal *et al.*, 2018). Fungal penetration attempts could be observed in all accessions (Supplementary Table S2). In more than 90% of the cases, host penetration occurred through the stomatal openings (Fig. 2d & d’).

### 3. Accelerated ROS accumulation and augmented cell death correlates with susceptibility

Typical host responses to pathogen invasion include a short oxidative burst followed by cell death. Therefore, we compared the early responses of the resistant and susceptible *Arabidopsis* accessions to *A. brassicae* with an emphasis on i) accumulation of H_2_O_2_ ii) spread of cell death.

During the early phases of infection, in a fraction of fungal penetration sites, the guard cells, adjoining epidermal cells and a small number of underlying mesophyll cells showed ROS accumulation suggestive of activation of host defence response. In the susceptible accessions the number of penetration sites associated with ROS production were marginally more than the resistant accessions. Quantitative analysis of the number of penetration sites associated with ROS accumulation as a factor of time revealed a greater increase in the number of such sites in the susceptible accessions as compared to the resistant accessions (Fig. 3c). Most significant difference was observed at 96 hpi where larger patches of DAB-stained cells were commonly observed at the spore inoculation site in the susceptible accessions while in the resistant accessions the expression of DAB was restricted to a few smaller and scattered patches (Fig. 3a, 3b, Supplementary Figure 1, and Table S3). The spread of ROS was measured by quantifying the area stained by DAB.

**Fig. 3.**
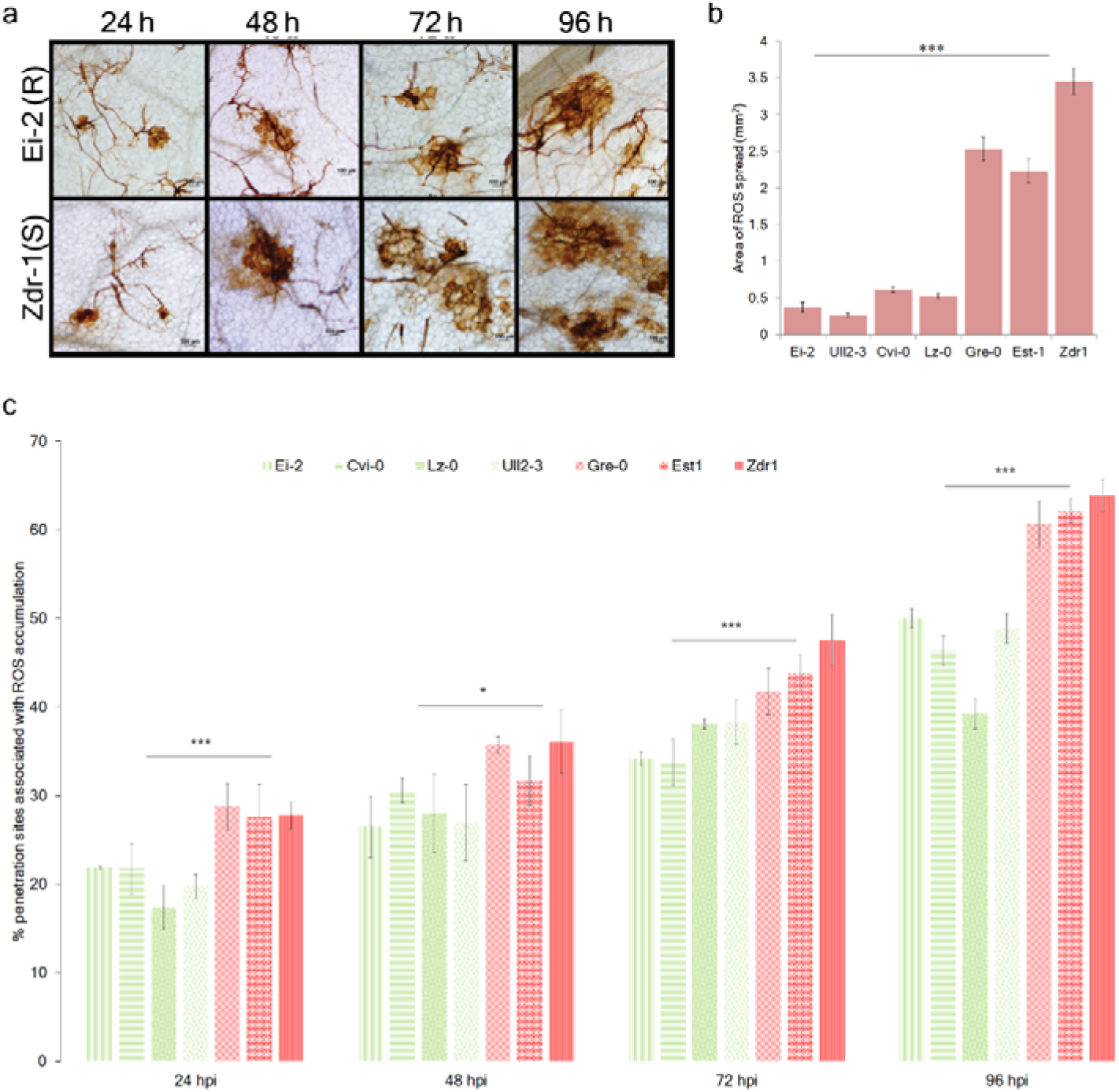
Differential H_2_O_2_ accumulations in resistant and susceptible *Arabidopsis* accessions upon *Alternaria brassicae* infection. (a) Representative microscopic images of H_2_O_2_ accumulation (brown color) in Ei-2 and Zdr-1 at 24, 48, 72 and 96 hpi. (b) Area of the spread of ROS production in response to *A. brassicae* infection in the contrasting subset of accessions at 96 hpi. For each accession, a total of 40 inoculation sites from three independent experiments were randomly selected and used for quantification of area of ROS spread. (c) Percentage of fungal penetration sites associated with H_2_O_2_ accumulation in leaves of the seven accessions infected with *A. brassicae*. Data is the mean of three biological replicates ± S.D. Significant differences were analysed with one-way ANOVA followed by Tukey’s HSD test at p < 0.05. * - p < 0.05; ** - p < 0.01; *** - p < 0.001

**Fig. 4.**
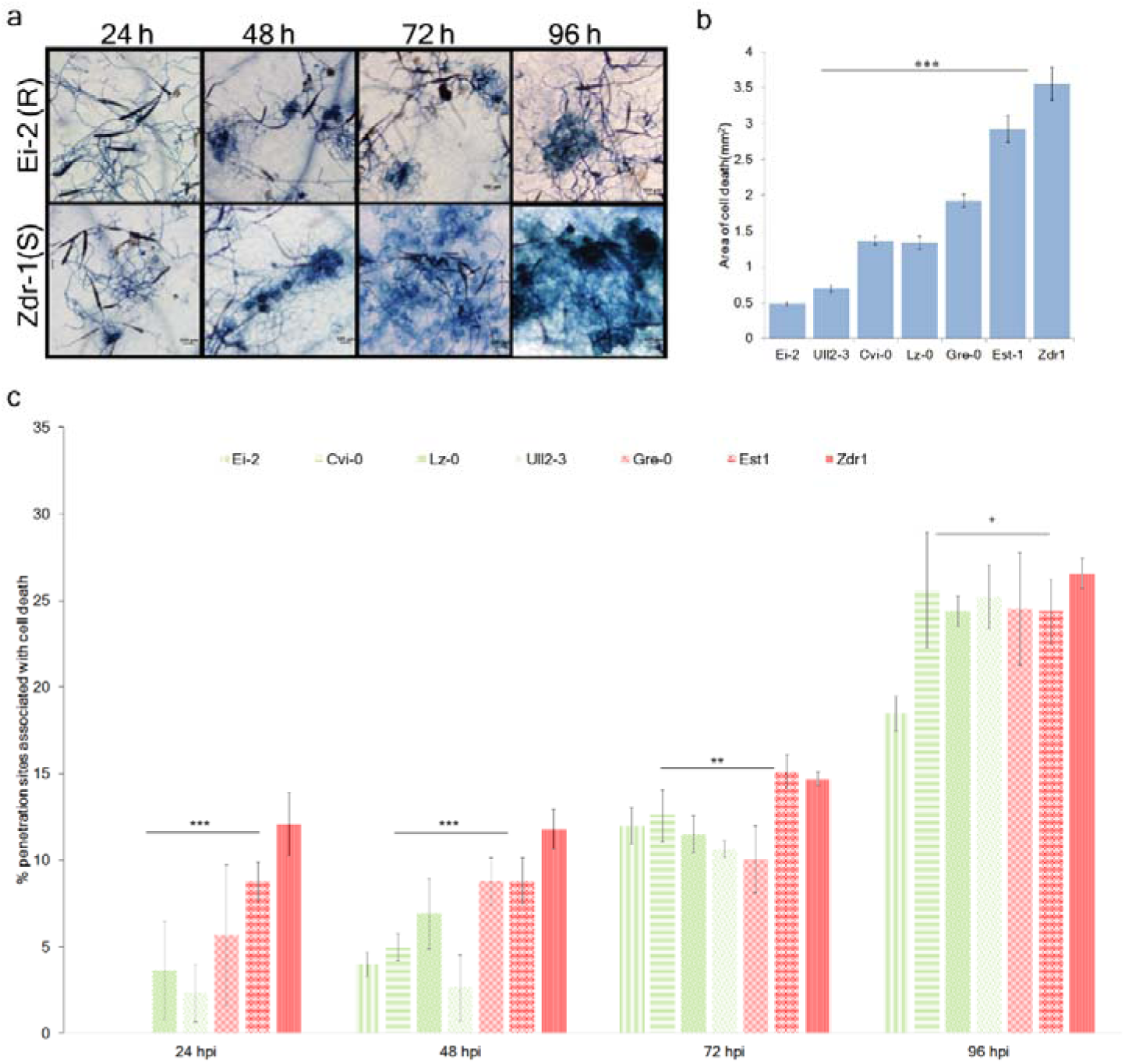
Cell death in resistant and susceptible *Arabidopsis* accessions upon *Alternaria brassicae* infection. (a) Representative microscopic images of cell death in the resistant accession (Ei-2) and susceptible accession (Zdr-1) at 24, 48, 72 and 96 hpi. (b) Area of spread of cell death in response to *A. brassicae* infection in the *Arabidopsis* accessions at 96 hpi. For each accession, a total of 40 inoculation sites from three independent experiments were randomly selected and used for quantification of area of cell death. (c) Percentage of *A. brassicae* penetration events associated with cell death in the resistant (Ull2-3, Lz-0, Cv8, Ei-2) and the susceptible (Zdr-1, Est-1, Gre3.2) *Arabidopsis* accessions. Data is the mean of three biological replicates ± S.D. Significant differences were analysed with one-way ANOVA followed by Tukey’s HSD test at p < 0.05. * - p < 0.05; ** - p < 0.01; *** - p < 0.001

Besides ROS, cell death was also observed at the inoculation site in the susceptible and resistant accessions. At the site of stomatal penetration (24 hpi), cell death was observed only in single epidermal cells, and by 48 hpi, the cell death was induced in the underlying mesophyll cells as well. A quantitative analysis of penetration-associated cell death was carried out temporally. During the initial time point, i.e. 24 hpi, cell death was seen in a small number of penetration sites in the susceptible accessions whereas none was observed in the resistant accessions. As time progressed, the number of penetration site associated with cell death also increased (Fig. 4a & c). Massive cell death was seen at 72 and 96 hpi as trypan blue stained, collapsed mesophyll cells (Fig. 4a; Supplementary Fig. 2). The spread of cell death as measured by the area of trypan blue staining was significantly different between the resistant and susceptible subsets (Fig. 4b, Supplementary Table S3).

### 4. Role of ROS in *Alternaria brassicae*-*Arabidopsis* interactions

To further investigate the role of ROS in the pathogenesis of *A. brassicae* we tested mutants of some of the known Respiratory Burst Oxidase Homologs (RBOHs) for their response to *A. brassicae* infection. Plasma membrane-bound NADPH oxidases (RBOHD and F) and cell wall peroxidases are considered as main sources of an oxidative burst in the apoplast (Sagi & Fluhr 2006). The homozygous T-DNA knockout mutant lines for *rbohD* (CS9555), *E* (SALK_146126C) and *F* (CS9557) showed highly reduced fungal establishment, as the lesions formed at 7dpi were fewer and smaller in size as compared to Col-0 (Fig. 5a; Supplementary Table S4). Interestingly, *rbohE* displayed the highest resistance of all the mutants tested (Fig. 5a). In addition to the reduced accumulation of H_2_O_2_, the mutants also displayed a reduced cell death compared with the wild type plants after inoculation with the *A. brassicae* spores (Fig. 5b & c; Supplementary Table S4).

**Figure 5.**
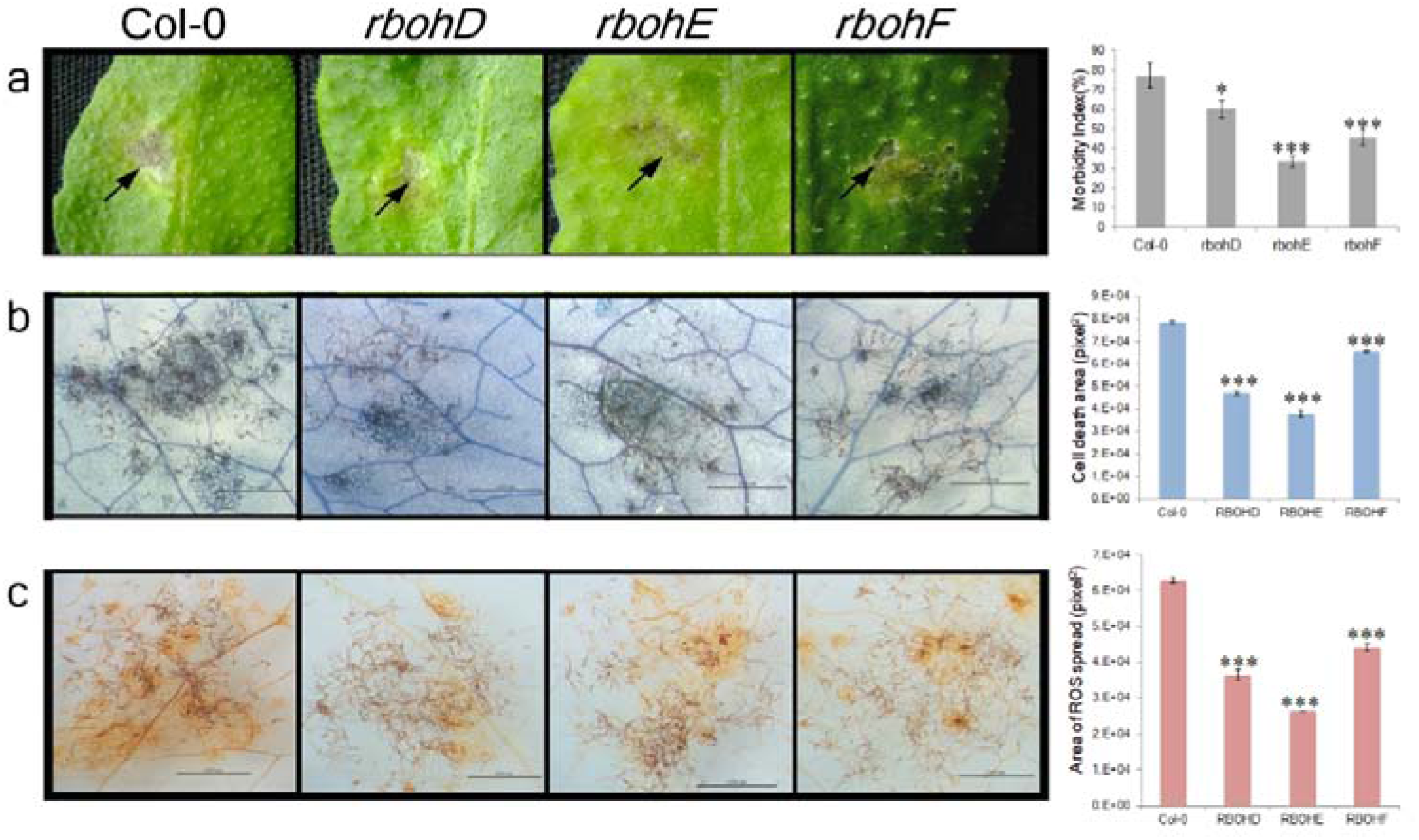
Phenotypic and cellular characterization of *Arabidopsis rbohD, E* and *F* plants. (a) Photographs of representative 5-6 week old mutants and WT (Col-0) plants showing the formation of lesion phenotype at 7 dpi. The graph represents the rate of disease incidence or morbidity (%) in the wild type and mutants. (b) Representative images of *A. brassicae* induced cell death (trypan blue staining) in the mutants and wild type. The graph represents the quantification of the area of induced cell death. (c) Representative images of DAB stained leaves of Col-0, *rbohD, rbohE* and *rbohF* plants to estimate H_2_O_2_ accumulation. The graph represents the area of H_2_O_2_ spread in the mutants and wild type. Trypan blue (cell death) and DAB (H_2_O_2_) stained areas were quantified by ImageJ. All experiments were repeated at least three times with at least 6 plants per experiment. Significant differences were analysed with unpaired t-test with unequal variances at p < 0.05. * - p < 0.05; ** - p < 0.01; *** - p < 0.001

To further analyse the role of RBOHs, we determined the expression levels of *RBOHD, E* and *F*, upon infection in both resistant (Ei-2) and susceptible (Zdr-1) accessions. During the interaction with *A. brassicae*, all the genes were induced, and their expression level was generally higher in the resistant accession as compared to the susceptible one at 48 hpi. *RBOHE* expression, however, exhibited a delayed upregulation with stronger expression at 4 dpi (Supplementary Fig. 3).

### 5. JA and ET signalling pathways contribute to resistance against *A. brassicae*

Apart from cytological analysis, to further dissect the resistance mechanism in the selected *Arabidopsis* accessions we analysed the expression profile of an array of defence signalling pathways upon *A. brassicae* infection. It is well established that pathogen invasion triggers the accumulation of several signalling molecules like salicylic acid (SA), jasmonic acid or ethylene (JA/ET) which in turn alter the expression of several defence genes. Here, we analysed the changes in expression for genes involved in either synthesis of the signalling molecules or those involved in downstream signalling in the two representative accessions, i.e. Ei-2 (resistant) and Zdr-1 (susceptible). Plants infected with *A. brassicae* or mock treated were sampled at2 and 4 dpi. Gene expression analysis using quantitative RT-PCR revealed that the JA and ET inducible marker, PDF1.2 was highly upregulated at 2 dpi in Ei-2 (Resistant) as compared to Zdr-1 (Susceptible). Whereas the SA responsive gene, PR1 was highly upregulated in Zdr-1 at 2 dpi as compared to Ei-2 (Fig. 6; Supplementary Table S6).

**Figure 6:**
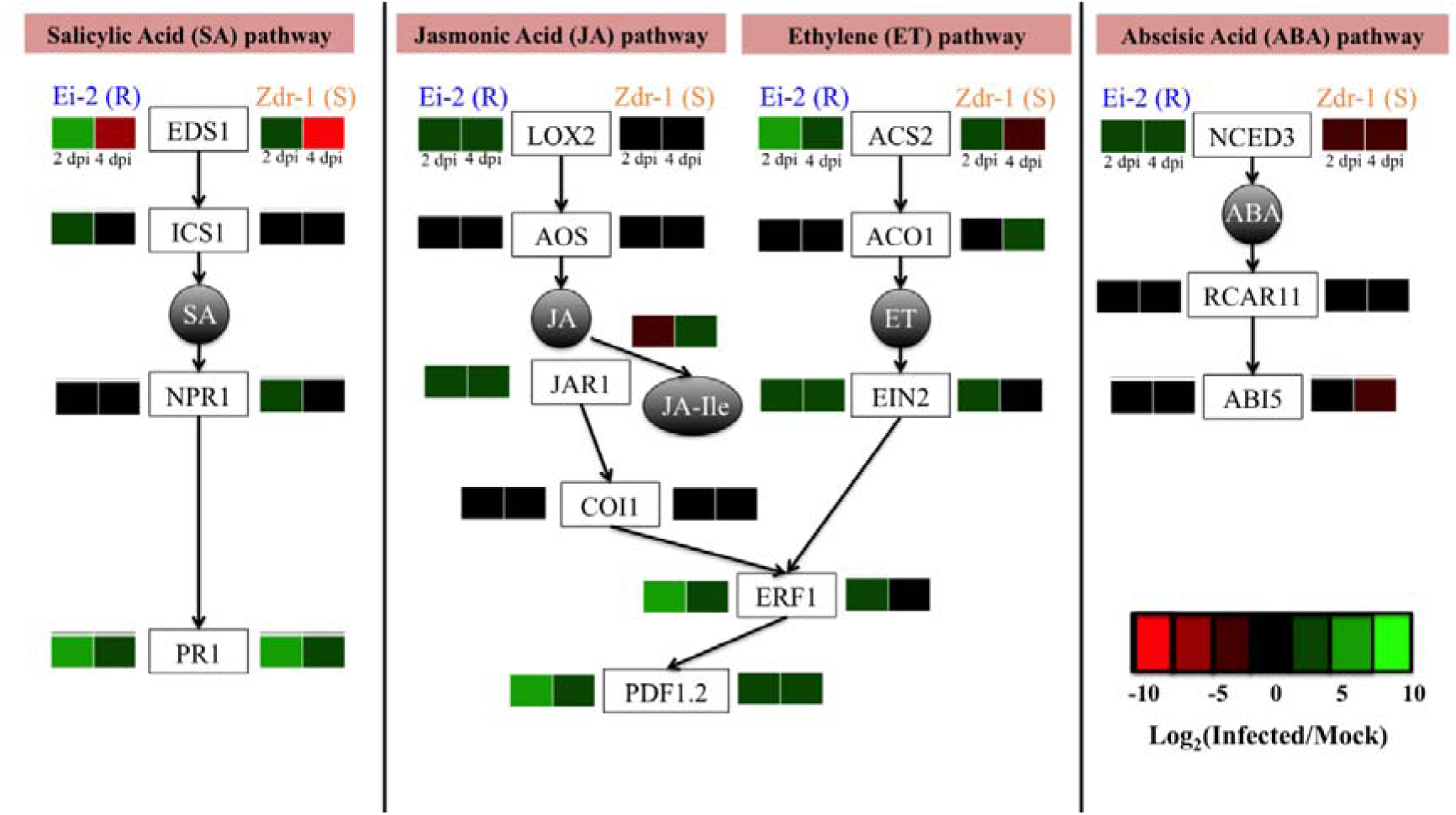
Summary of defence gene expression in susceptible and resistant accession of *Arabidopsis thaliana* after inoculation with *Alternaria brassicae*. Leaves of 5-6 week-old plants of Ei-2 (resistant) and Zdr-1 (susceptible) were inoculated with *A. brassicae* and harvested at different times after inoculation (2 and 4 dpi). The expression values of various genes were determined by qPCR and represented as heatmaps overlaid onto the different phytohormonal pathways. Details of expression analysis are given in Material and Methods and the fold change values are given in Supplementary Table S6

Additionally, LOX2 and ACS2 genes, which are involved in the JA and ET biosynthesis pathways, displayed a stronger upregulation in the resistant accession as compared to the susceptible accession. This is consistent with previous reports, which have observed a major positive contribution of JA and ET pathways in defence against necrotrophs (Glazebrook 2005; Thomma *et al.*, 1998; Tsuda *et al.*, 2009). However, we did not observe any significant antagonism of the SA pathway by JA in the resistant accession, Ei-2. The SA pathway genes such as EDS1 and ICS1 were significantly upregulated in both the accessions at 2 dpi but the quantum of upregulation was higher in the resistant one. Nevertheless, PR1 was significantly more upregulated in the susceptible accession as compared to the resistant accession at both 2 and 4 dpi. ABA, another phytohormone, has been attributed positive and negative regulatory functions depending on the pathosystem analysed. We analysed the expression levels of three known genes in the ABA pathway viz. NCED3, RCAR11 and ABI5. NCED3, associated with ABA synthesis was upregulated strongly by 4 dpi in the resistant accession Ei-2, and was downregulated in the susceptible accession Zdr-1 (Fig. 6; Supplementary Table S6).

## Discussion

Our previous study investigating the interaction between *Arabidopsis* and *Alternaria brassicae* suggests the involvement of multiple mechanisms conferring resistance against the necrotrophic fungus (Rajarammohan *et al.*, 2018). The differential response of *Arabidopsis* accessions to *A. brassicae* infection provides an excellent handle to address some of the fundamental questions in *A. brassicae* host interactions. The main goal of this work was to study the cellular and molecular intricacies of *Arabidopsis thaliana* -*A. brassicae* interaction. Thus, a total of seven *Arabidopsis* accessions originating from different geographic origins, responding differentially to *A. brassicae* infection were selected for a comparative histological and molecular analysis.

Preformed defences consisting of epicuticular waxes, antimicrobial compounds on plant surface may contribute to the differential phenotypes observed in response to *A. brassicae* infection. A positive correlation between the epicuticular waxes and resistance against *A. brassicae* in Brassicas has been reported earlier (Saharan *et al.*, 2016). Epicuticular waxes reduce spore adhesion and spore germination in the resistant cultivars. Moreover, stomatal aperture and number of stomata appear to be contributing factors to the tolerance of *A. brassicae* in Brassicas. In *Arabidopsis*-*A. brassicae* interaction, no distinctive differences in the initial spore germination, germ-tube formation, and penetration were observed between the resistant and susceptible accessions suggesting that the basis of enhanced tolerance of Ei-2, Ull2-3, Cv8 and Lz-0 accessions does not reside in the early pre-penetration stages of host-pathogen interaction.

Generation of ROS has been shown to be facilitating pathogenesis of necrotrophs. In our study, we observed a small proportion of penetration sites associated with H_2_O_2_ accumulation in both resistant and susceptible accessions. As the infection progressed, massive accumulation of H_2_O_2_ was observed in large patches of epidermal as well as mesophyll cells in the susceptible accessions. Similar patterns were observed for the associated cell death in resistant and susceptible accessions. Therefore, we postulated that the induction of ROS and the associated cell death induced by *A. brassicae* might be essential for its establishment and may require host NADPH oxidases. The *Arabidopsis* genome encodes at least 10 RBOH genes of which RBOHD, E, and F have been reported to be pathogen-inducible. Analysis of *rbohD, E* and *F* mutants and wild-type (Col-0) revealed that the mutants produced lesser amounts of ROS upon pathogen challenge. The mutants displayed lower morbidity (*rbohE>rbohF>rbohD>*Col-0) and also had smaller patches of dead cells as compared to wild type (Col-0) (Fig. 5). This is in concordance with the recent study by Ranjan *et al.*, 2017, which showed that the pathogenic invasion of *S. sclerotiorum* requires the host RBOHs and the silencing of a specific RBOH in soybean led to enhanced resistance against the pathogen. In addition to host generated ROS, *A. brassicae* is also known to produce ROS at the penetrating hyphal tips (Mandal *et al.*, 2018). The role of pathogen-produced ROS in the *Arabidopsis-A. brassicae* pathosystem needs to be addressed, given the various studies that show the importance of pathogen-produced ROS in development of infection structures and host penetration (Foley *et al.*, 2016; Samalova *et al.*, 2014; Segmuller *et al.*, 2008; Shetty *et al.*, 2007; Tiedemann 1997).

In order to understand the signalling mechanisms underlying defence against *A. brassicae*, we examined the expression profiles of genes belonging to various phytohormonal signalling pathways in response to infection. The phytohormones salicylic acid (SA), jasmonic acid (JA), ethylene (ET) comprise a complex set of hormone signalling pathways that lead to the regulation of resistance in plants against necrotrophic pathogens. We examined genes relating to biosynthesis, signalling and downstream targets of each of the phytohormones. We found that the JA and ET pathways were strongly upregulated in the resistant accession as compared to the susceptible one. This is in line with previous reports of plant defence against necrotrophic pathogens wherein resistance is conferred by JA/ET signalling pathways (Glazebrook J 2005; Mengiste 2012; Thomma *et al.*, 2001; van Kan 2006). Abscisic acid (ABA) is known to inhibit the pathogen-mediated SA response (Mazumder *et al.*, 2013) and NCED3, a key biosynthetic enzyme in the ABA pathway was strongly upregulated in the resistant accession Ei-2, and was downregulated in the susceptible accession Zdr-1. Also, it is well established that ABA induces closure of stomata. Stomata have been shown to be the major penetration site of *A. brassicae* in our previous study (Mandal *et al.*, 2018). Therefore, ABA might have an important role in defence against *A. brassicae*. ROS generated by the NADPH oxidases is known to act both upstream and downstream of the SA signalling pathway and levels of SA modulate ROS homeostasis, whereas JA is known to be involved in inhibiting ROS production at later stages and thus contributing to lesion containment (Overmyer *et al.*, 2003). Though the current study shows a definitive link between ROS, associated cell death and susceptibility to *A. brassicae*, the exact mechanism of how ROS induces cell death is not known. Detailed genetic analysis of the interaction of ROS and the phytohormonal pathways would lead to a deeper understanding of the mechanism of ROS mediated susceptibility.

In conclusion, this study reveals the central role of ROS and cell death in the *Arabidopsis-A. brassicae* interaction. The increased accumulation of ROS and associated cell death led to an increase in susceptibility. Expression profiling of hormonal signalling pathways suggests a positive role for JA, ET, and ABA in resistance to *A. brassicae*. The interaction of ROS with the phytohormonal pathways and the balance between them may serve to fine-tune the defence response against *A. brassicae.*

## Contributions

JK conceived and designed research. SM performed the microscopic analysis of the accessions and mutant lines and analysed the data. SR performed the gene expression studies, statistical analysis, and contributed to the interpretation of the data. JK, SM, and SR wrote the manuscript. All authors read and approved the manuscript.

## Acknowledgements

This work was financially supported by the grants from Science and Engineering Research Board (SB/FT/LS-327/2012) and Delhi University-Department of Science and Technology PURSE Grant (RC/2016/744). Research fellowship to SM and SR from University Grants Commission (UGC), Government of India is acknowledged. We also acknowledge Central Instrument Facility-University of Delhi, South Campus and Centre for Genetically Modified Crop Plants (CGMCP), University of Delhi for sharing facilities and plant growth space. The authors declare that they have no conflict of interest.

## Supplementary Information Legends

**Table S1** *A. brassicae* spore germination (24 hpi) on leaf surface of *Arabidopsis* accessions

**Table S2** Penetration frequency of *Alternaria brassicae* on different *Arabidopsis* accessions at different hours post inoculation (hpi)

**Table S3** F-values and p-values from a single factor ANOVA to determine significant differences between the accessions in the host responses viz. H_2_O_2_ production and cell death

**Table S4** p-values from unpaired t-test with unequal variances to determine significant differences between *rboh* mutants and Col-0 (wild type) in the host responses viz. morbidity, H_2_O_2_ production and cell death

**Table S5** List of primers used in expression profiling of hormonal pathway genes

**Table S6** Expression values in terms of Fold Change (FC) of the genes of various hormonal pathways in Ei-2 and Zdr1 upon infection.

**Supplementary Figure 1** Microscopic detection of ROS by DAB at 24, 48, 72 and 96 hpi shows enhanced accumulation of ROS in susceptible accessions (Est-1, Ull2-3) in comparison to resistant accessions (Lz-0, Cv8)

**Supplementary Figure 2** Trypan blue staining for detection of cell death in susceptible and resistant accessions at 24, 48, 72 and 96 hpi with *A. brassicae*. Representative pictures show the spread of cell death is more accelerated in the susceptible accessions (Est-1 and Ull2-3) as compared to the resistant accessions (Lz-0 and Cv8), where the spread is confined. DC: Dead cells, S: Spore, H: Hyphae

**Supplementary Figure 3** Expression patterns of RBOHD, E, and F in Ei-2 and Zdr-1, 2 and 4 days post infection (dpi) w.r.t. mock infected (distilled water). The mean values (±SD) of three biological replicates are shown. Expression levels were normalised to the expression values of endogenous control-TIP41-like gene (At4g34270)

**Supplementary Figure 4** Confirmation of T-DNA insertion in *rbohD, E*, and *F* mutants. The position of the T-DNA insertions in the genes are shown along with the gel images of amplification with insertion specific primers in Col-0 (wild type) and the mutant lines.

## References

1. Bartsch M, Gobbato E, Bednarek P, et al., 2006. Salicylic acid-independent ENHANCED DISEASE SUSCEPTIBILITY1 signaling in Arabidopsis immunity and cell death is regulated by the monooxygenase FMO1 and the Nudix hydrolase NUDT7. The Plant Cell 18, 1038–51.

2. Ferrari S, Plotnikova JM, De Lorenzo G and Ausubel FM, 2003. Arabidopsis local resistance to Botrytis cinerea involves salicylic acid and camalexin and requires EDS4 and PAD2, but not SID2, EDS5 or PAD4. The Plant Journal 35, 193–205.

3. Foley RC, Kidd BN, Hane JK, Anderson JP and Singh KB, 2016. Reactive Oxygen Species Play a Role in the Infection of the Necrotrophic Fungi, Rhizoctonia solani in Wheat. PLoS One 11, e0152548.

4. Glawischnig E, Hansen BG, Olsen CE and Halkier BA, 2004. Camalexin is synthesized from indole-3-acetaldoxime, a key branching point between primary and secondary metabolism in Arabidopsis. Proceedings of the National Academy of Sciences of the United States of America 101, 8245–50.

5. Glazebrook J, 2005. Contrasting mechanisms of defense against biotrophic and necrotrophic pathogens. Annual Review of Phytopathology 43, 205–27.

6. Govrin EM and Levine A, 2000. The hypersensitive response facilitates plant infection by the necrotrophic pathogen Botrytis cinerea. Current Biology 10, 751–7.

7. Govrin EM and Levine A, 2002. Infection of Arabidopsis with a necrotrophic pathogen, Botrytis cinerea, elicits various defense responses but does not induce systemic acquired resistance (SAR). Plant Molocular Biology 48, 267–76.

8. Mandal S, Rajarammohan S and Kaur J, 2018. Alternaria brassicae interactions with the model Brassicaceae member Arabidopsis thaliana closely resembles those with Mustard (Brassica juncea). Physiological and Molecular Biology of Plant 24, 51–9.

9. Marino D, Dunand C, Puppo A and Pauly N, 2012. A burst of plant NADPH oxidases. Trends in Plant Science 17, 9–15.

10. Mazumder M, Das S, Saha U, Chatterjee M, Bannerjee K and Basu D, 2013. Salicylic acid-mediated establishment of the compatibility between Alternaria brassicicola and Brassica juncea is mitigated by abscisic acid in Sinapis alba. Plant Physiology Biochemistry 70, 43–51.

11. Mengiste T, 2012. Plant immunity to necrotrophs. Annual Review of Phytopathology 50, 267–94.

12. Overmyer K, Brosche M and Kangasjarvi J, 2003. Reactive oxygen species and hormonal control of cell death. Trends in Plant Science 8, 335–42.

13. Rajarammohan S, Kumar A, Gupta V, Pental D, Pradhan AK and Kaur J, 2017. Genetic Architecture of Resistance to Alternaria brassicae in Arabidopsis thaliana: QTL Mapping Reveals Two Major Resistance-Conferring Loci. Frontiers in Plant Science 8, 260.

14. Rajarammohan S, Pradhan Ak, Pental D and Kaur J, 2018. Genome-wide association mapping in Arabidopsis identifies novel genes underlying quantitative disease resistance to Alternaria brassicae. Molecular Plant Pathology 19, 1719–32.

15. Ranjan A, Jayaraman D, Grau C, et al, 2017. The pathogenic development of Sclerotinia sclerotiorum in soybean requires specific host NADPH oxidases. Molecular Plant Pathology 19, 700–14.

16. Rate DN and Greenberg JT, 2001. The Arabidopsis aberrant growth and death2 mutant shows resistance to Pseudomonas syringae and reveals a role for NPR1 in suppressing hypersensitive cell death. The Plant Journal 27, 203–11.

17. Sagi M and Fluhr R, 2006. Production of reactive oxygen species by plant NADPH oxidases. Plant Physiology 141, 336–40.

18. Saharan GS, Mehta N and Meena PD, 2016. Alternaria Diseases of Crucifers: Biology, Ecology and Disease Management: Springer.

19. Samalova M, Meyer AJ, Gurr SJ and Fricker MD, 2014. Robust anti-oxidant defences in the rice blast fungus Magnaporthe oryzae confer tolerance to the host oxidative burst. New Phytologist 201, 556–73.

20. Segmuller N, Kokkelink L, Giesbert S, Odinius D, Van Kan J and Tudzynski P, 2008. NADPH oxidases are involved in differentiation and pathogenicity in Botrytis cinerea. Molecular Plant-Microbe Interactions 21, 808–19.

21. Shetty NP, Jørgensen HJL, Jensen JD, Collinge DB, Shetty HS, 2008. Roles of reactive oxygen species in interactions between plants and pathogens. European Journal of Plant Pathology 121, 13.

22. Shetty NP, Mehrabi R, Lutken H, et al, 2007. Role of hydrogen peroxide during the interaction between the hemibiotrophic fungal pathogen Septoria tritici and wheat. New Phytologist 174, 637–47.

23. Broekaert WF Thomma BP, Eggermont K, Penninckx IA, et al, 1998. Separate jasmonate-dependent and salicylate-dependent defense-response pathways in Arabidopsis are essential for resistance to distinct microbial pathogens. Proceedings of the National Academy of Sciences of the United States of America 95, 15107–11.

24. Thomma BP, Penninckx IA, Broekaert WF, 2001. The complexity of disease signaling in Arabidopsis. Current Opinion in Immunology 13, 63–8.

25. Thordal-Christensen H, Zhang Z, Wei Y and Collinge DB, 1997. Subcellular localization of H2O2 in plants, H2O2 accumulation in papillae and hypersensitive response during barley-powdery mildew interaction. The Plant Journal 11, 1187–94.

26. Tiedemann AV, 1997. Evidence for a primary role of active oxygen species in induction of host cell death during infection of bean leaves with Botrytis cinerea. Physiological and Molecular Plant Pathology 50, 151–66.

27. Tsuda K, Sato M, Stoddard T, Glazebrook J and Katagiri F, 2009. Network properties of robust immunity in plants. PLoS Genetics 5, e1000772.

28. van Baarlen P, van Belkum A, Summerbell RC, Crous PW, Thomma BP, 2007. Molecular mechanisms of pathogenicity: how do pathogenic microorganisms develop cross-kingdom host jumps? FEMS Microbiology Reviews 31, 239–77.

29. Van Breusegem F, Dat JF, 2006. Reactive oxygen species in plant cell death. Plant Physiology 141, 384–90.

30. van Kan JA, 2006. Licensed to kill: the lifestyle of a necrotrophic plant pathogen. Trends in Plant Science 11, 247–53.

31. Willems E, Leyns L, Vandesompele J, 2008. Standardization of real-time PCR gene expression data from independent biological replicates. Analytical Biochemistry 379, 127–9.

32. Williams B, Kabbage M, Kim HJ, Britt R, Dickman MB, 2011. Tipping the balance: Sclerotinia sclerotiorum secreted oxalic acid suppresses host defenses by manipulating the host redox environment. PLOS Pathogens 7, e1002107.

33. Wolpert TJ, Dunkle LD, Ciuffetti LM, 2002. Host-selective toxins and avirulence determinants: what’s in a name? Annual Review of Phytopathology 40, 251–85.

