## Supplementary Data for "ROS accumulation and associated cell death mediates susceptibility to *Alternaria brassicae* in *Arabidopsis* accessions"

### **Supplementary Tables and Figures**

**Table S1:** *A. brassicae* spore germination (24 hpi) on leaf surface of *Arabidopsis* accessions

| S.No. | Accessions | Interaction Phenotype | Total no. of spores* | Total no. of germinated spores | (%) Germination |
|---|---|---|---|---|---|
| 1. | Zdr-1 | Susceptible | 2906 | 2894 | 99.59 |
| 2. | Est-1 | Susceptible | 2870 | 2860 | 99.65 |
| 3. | Ull2-3 | Resistant | 2799 | 2792 | 99.75 |
| 4. | Lz-0 | Resistant | 2886 | 2876 | 99.65 |
| 5. | Cv8 | Resistant | 2891 | 2881 | 99.65 |
| 6. | Ei-2 | Resistant | 2864 | 2856 | 99.72 |

* Total number of spore represents spore count from all the host responses ($H_2O_2$, cell death and callose deposition). The experiment was repeated three times with similar results.

**Table S2:** Penetration frequency of *Alternaria brassicae* on different *Arabidopsis* accessions at different hours post inoculation (hpi)

| Accessions | 24 hpi | 48 hpi | 72 hpi | 96 hpi |
|---|---|---|---|---|
| | Total penetration site● | Total penetration site● | Total penetration site● | Total penetration site● |
| **Zdr-1** | 365 | 1018 | 1786 | 1894 |
| **Est-1** | 338 | 952 | 1720 | 1839 |
| **Ull2-3** | 331 | 795 | 1598 | 1671 |
| **Lz-0** | 341 | 731 | 1558 | 1709 |
| **Cv8** | 343 | 972 | 1685 | 1709 |
| **Ei-2** | 329 | 805 | 1556 | 1761 |

- Total penetration sites represent the sum of all the penetration sites observed in experiments evaluating $H_2O_2$ production, cell death and callose deposition as a result of host responses upon infection.

**Table S3**: F-values and p-values from a single factor ANOVA to determine significant differences between the accessions in the host responses viz. $H_2O_2$ production and cell death

| **$H_2O_2$ production** | **24 hpi** | **48 hpi** | **72 hpi** | **96 hpi** |
|---|---|---|---|---|
| F-value | 15.84 | 4.00 | 11.52 | 53.14 |
| p-value | 4.54E-05 | 0.01966 | 0.0002236 | 5.99E-08 |
| **Cell Death** | | | | |
| F-value | 17.86 | 13.94 | 5.54 | 4.19 |
| p-value | 2.44E-05 | 8.71E-05 | 0.005833 | 0.01675 |
| **Area of $H_2O_2$ spread** | **96 hpi** | | | |
| F-value | 126.07 | | | |
| p-value | <2.00E-16 | | | |
| **Area of Cell death** | **96 hpi** | | | |
| F-value | 82.66 | | | |
| p-value | <2.00E-16 | | | |

**Table S4**: p-values from unpaired t-test with unequal variances to determine significant differences between *rboh* mutants and Col-0 (wild type) in the host responses viz. morbidity, $H_2O_2$ production and cell death

| | **Morbidity** | **Area of $H_2O_2$ spread** | **Area of cell death** |
|---|---|---|---|
| *rbohD* | 0.023441706 | 0.000170902 | 2.86979E-06 |
| *rbohE* | 0.00224845 | 7.26745E-05 | 7.41458E-05 |
| *rbohF* | 0.003333658 | 0.000105114 | 7.73834E-06 |

**Table S5:** List of primers used in expression profiling of hormonal pathway genes

| S.No. | Primer Name | Primer Sequence |
|---|---|---|
| 1 | RTJAR1F | GGAAGGAGAGGAGAAACCG |
| 2 | RTJAR1R | CGATACAACCCCTGCGTAAT |
| 3 | RTLOX2F | GCGGATCTCATCAAAAGGG |
| 4 | RTLOX2R | TCACATAGTCTGTCACCCATTCTT |
| 5 | RTCOI1F | GTTCTCGAGGACAAGGAATGT |
| 6 | RTCOI1R | TAAGCCTTCTTCGTCCTCCA |
| 7 | RTERF1F | AGTCAAGACCTTCCGATCAA |
| 8 | RTERF1R | CGAGCCAAACCCTAATACC |
| 9 | RTACS2F | TATGCCGCATTTGATAGAGAC |
| 10 | RTACS2R | AGTGGATTTGATGGGTTGGT |
| 11 | RTACO1F | CGACGATTACAGAACGTTAATGA |
| 12 | RTACO1R | GTGGATAATTGCTGACTTTGGTT |
| 13 | RTEIN2F | GCCTACAGGGAGTGATTGA |
| 14 | RTEIN2R | GGTTGTGCATTTGCCTTT |
| 15 | RTICS1F | CTAGTCGAAAAGGGTCTTGG |
| 16 | RTICS1R | GATCAAGCTCATTCCACTCTG |
| 17 | RTEDS1F | ATTCAGGTGATCGAGCAAGA |
| 18 | RTEDS1R | AACCAAGCACCTCGTCAA |
| 19 | RTNPR1F | TTGAAAATAGAGTTGCACTTGC |
| 20 | RTNPR1R | TACCAGTGAGACGGTCAGG |
| 21 | RTPR1F | TAGCCCACAAGATTATCTAA |
| 22 | RTPR1R | CACCAGAGTGTATGAGTC |
| 23 | RS_At3G14440F | AGTAAACCGGCAAGCGTAG |
| 24 | RS_At3G14440R | TTTGCTCCGGTGAATGAA |
| 25 | RS_At2G36270F | ACTGTATATGCTTGTTTTCTTGCT |
| 26 | RS_At2G36270R | GACATGGACAAGTGGATAACA |
| 27 | RS_At4G17870F | GATTACCGGCGAACACAT |
| 28 | RS_At4G17870R | GTCCAGATCCGATTCTCTTT |
| 29 | RTPDF1.2F | TTGCTGCTTTCGACGCA |
| 30 | RTPDF1.2R | TGTCCCACTTGGCTTCTCG |

**Table S6:** Expression values in terms of Fold Change (FC) of the genes of various hormonal pathways in Ei-2 and Zdr1 upon infection. The $Log_2$(FC) values of each gene for the corresponding samples are given.

| | **E2** | **E4** | **Z2** | **Z4** |
|---|---|---|---|---|
| ***LOX2*** | 3.23603155 | 1.98365072 | 1.519546 | -0.5370375 |
| ***AOS*** | 1.15969247 | -0.8749296 | 0.17322767 | -0.3754309 |
| ***JAR1*** | 2.15559446 | 1.97311411 | -1.8048997 | 1.96484427 |
| ***COI1*** | 0.43115365 | -0.4092358 | -0.2166549 | -0.146533 |
| ***ERF1*** | 4.92337706 | 2.23287188 | 2.69416917 | 0.33822291 |
| ***PDF1.2*** | 6.79665313 | 3.90847788 | 4.10957826 | 2.17117961 |
| ***EDS1*** | 6.24441895 | -6.3293459 | 2.83414475 | -11.090252 |
| ***ICS1*** | 3.49466008 | -0.8484784 | 1.15603135 | -0.8196233 |
| ***NPR1*** | 1.32474717 | 0.6641304 | 1.75444457 | 0.21900243 |
| ***PR1*** | 4.94998489 | 2.92279018 | 6.02399743 | 3.52941831 |
| ***ACS2*** | 6.29899745 | 1.92312597 | 4.57989436 | -4.2125456 |
| ***ACO1*** | 1.34687291 | 1.15733223 | 0 | 1.64363604 |
| ***EIN2*** | 1.67160211 | 2.55874855 | 3.19635553 | 1.01246679 |
| ***NCED3*** | 1.65493761 | 4.54149499 | -3.8933637 | -3.5399053 |
| ***ABI5*** | 1.01030383 | 0.84623406 | 0.1919494 | -1.8820081 |
| ***RCAR11*** | 0.24290194 | 1.1059302 | -0.4267867 | 0.71463712 |

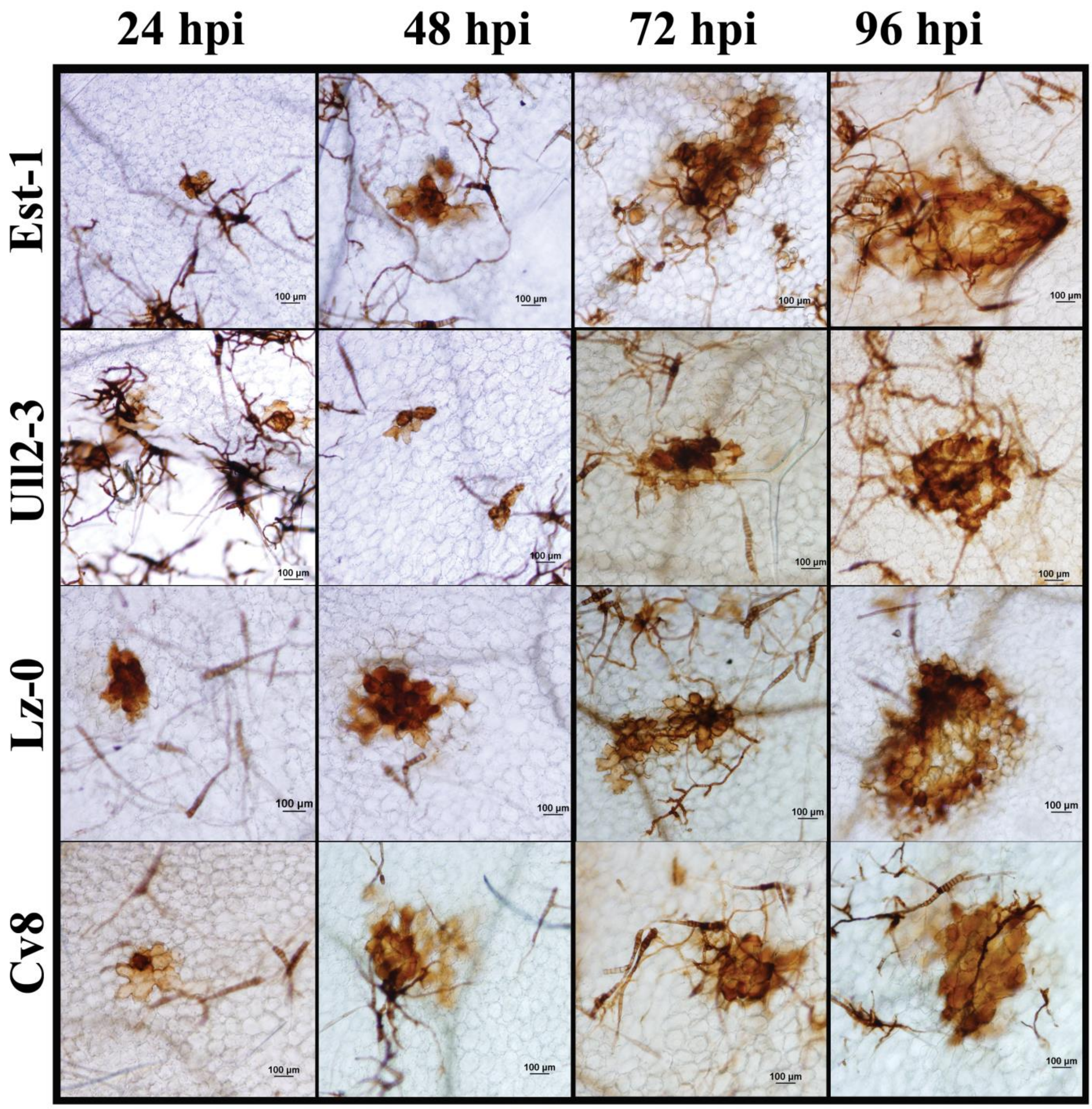


**Supplementary Figure 1:** Microscopic detection of ROS by DAB at 24, 48, 72 and 96 hpi shows enhanced accumulation of ROS in susceptible accessions (Est-1, Ull2-3) in comparison to resistant accessions (Lz-0, Cv8).

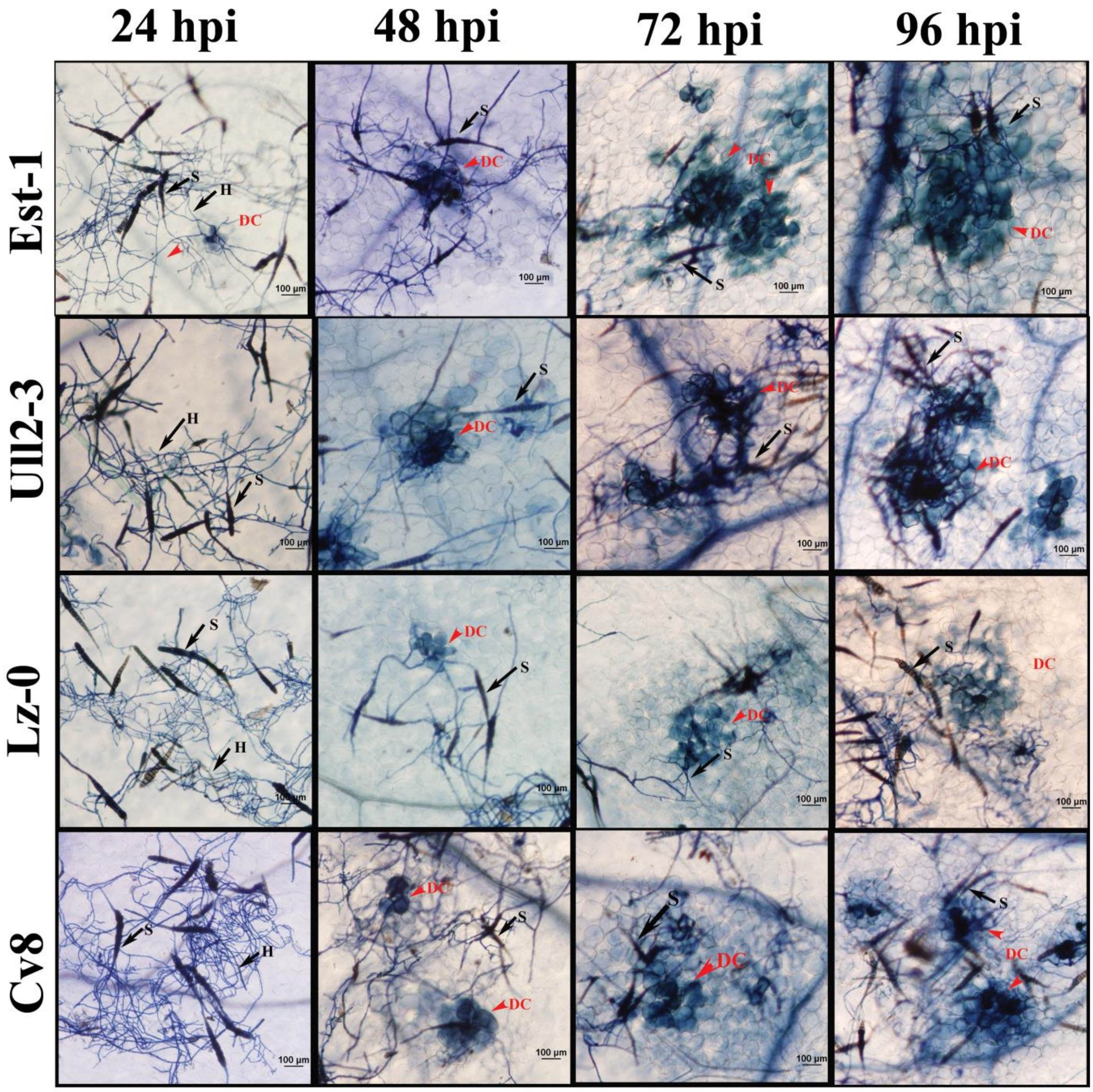


**Supplementary Figure 2:** Trypan blue staining for detection of cell death in susceptible and resistant accessions at 24-, 48-, 72- and 96- hpi with *A. brassicae*. Representative pictures show the spread of cell death is more accelerated in the susceptible accessions (Est1 and Ull2-3) as compared to the resistant accessions (Lz-0 and Cv8), where the spread is confined. DC= Dead cells, S: Spore, H: Hyphae.

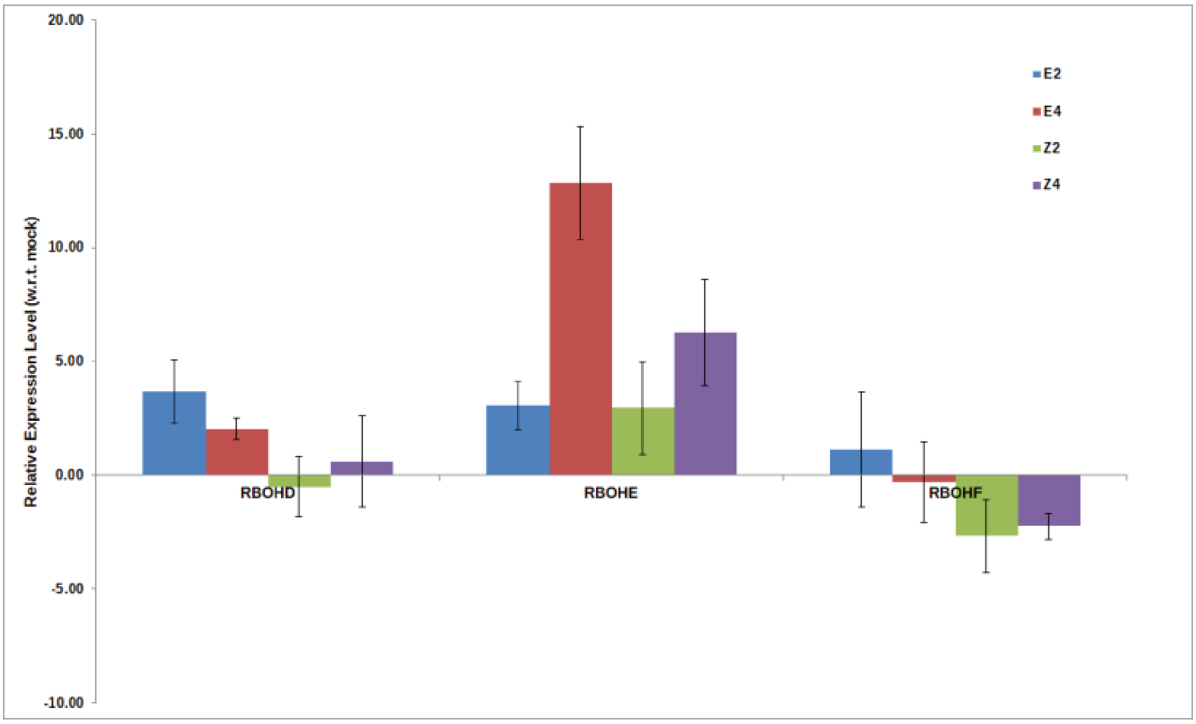


**Supplementary Figure 3:** Expression pattern of RBOHD, E, and F in Ei-2 and Zdr1, 2- and 4-days post infection (dpi) w.r.t. mock infected (distilled water). The mean values (±SD) of three biological replicates are shown. Expression levels were normalized to the expression values of endogenous control – TIP41-like gene (At4g34270)

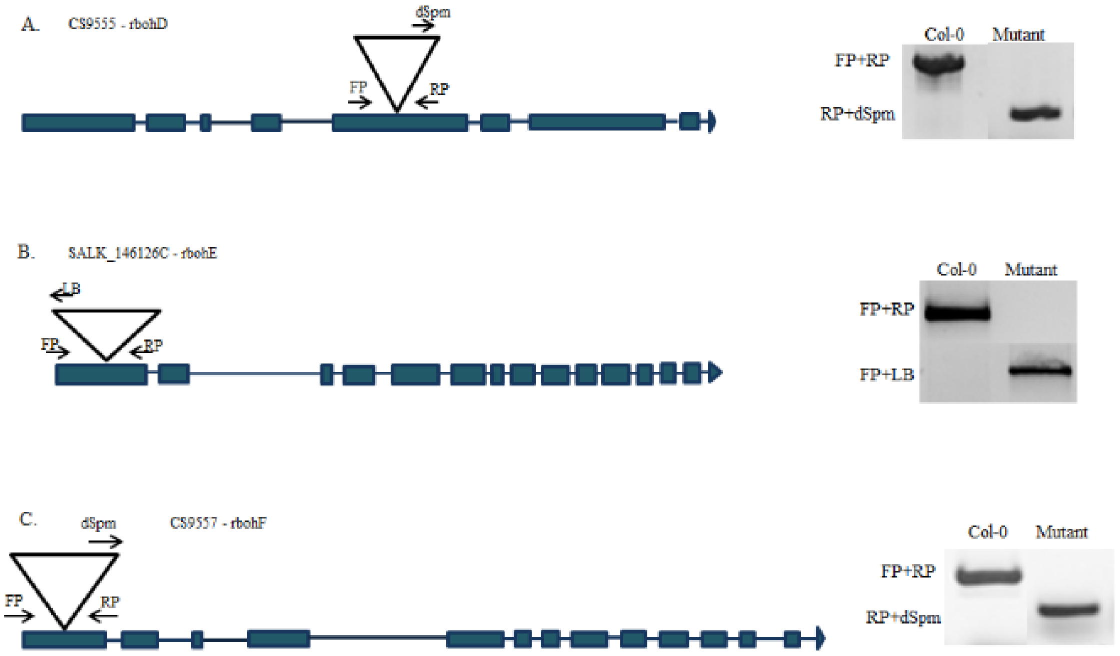


**Supplementary Figure 4:** Confirmation of T-DNA insertion in *rbohD*, *E*, and *F* mutants. The position of the insertions in the genes are shown along with the gel images of amplification with insertion specific primers in Col-0 (wild type) and the mutant lines.
